# Repair outcomes after germline homing endonuclease cleavage in Anopheles gambiae inform the design of synthetic gene drives

**DOI:** 10.64898/2026.06.23.733901

**Authors:** Daniel Naujoks, Tony Nolan

## Abstract

Homing endonuclease genes spread by cleaving homologous chromosomes that lack the endonuclease cassette, after which repair from the endonuclease-containing chromosome converts the cut allele into a copy of the drive allele. This mechanism has provided a conceptual foundation for synthetic gene drive systems, including CRISPR-based drives, that represent promising strategies for the genetic control of insect pests. However gene drive performance depends critically on the repair pathways available in the germline of the target organism. Here, we report a set of transgenic assays originally developed as part of an attempt to establish gene targeting in the malaria mosquito *Anopheles gambiae* using an in vivo-generated linear targeting molecule. Although the intended FLP-mediated excision step was not achieved in the mosquito germline, analysis of the component strains revealed efficient germline activity of the rare-cutting homing endonuclease I-SceI and a striking bias towards homology-based repair of I-SceI-induced double-strand breaks. Across reporter and donor configurations, cleavage outcomes were dominated by single-strand annealing, microhomology-mediated repair, synthesis-dependent strand annealing and gene conversion-like events, with comparatively limited evidence for classical non-homologous end joining. In reciprocal crosses designed to distinguish gene conversion from gamete loss, I-SceI cleavage also produced inheritance distortion consistent with both conversion of the cleaved allele and reduced recovery of gametes carrying extensively damaged donor alleles. These findings indicate that the *An. gambiae* germline can strongly favour homology-dependent repair following homing endonuclease cleavage and that cleavage can also generate meiotic drive-like distortion through selective loss of damaged gametes. The results have direct relevance for the design and interpretation of homing endonuclease and CRISPR-based gene drives in malaria mosquitoes, where the balance between homology-directed repair, end joining and gamete viability will determine drive efficiency, resistance formation and transmission bias.

**Author summary:** Gene drives depend on a simple but demanding principle: a nuclease cuts one chromosome, and the cell repairs the break using the homologous chromosome as a template, copying the drive element in the process. Before CRISPR, this type of system was explored using naturally occurring homing endonucleases such as I-SceI. We attempted to develop a gene targeting system in *Anopheles gambiae* based on the Rong and Golic strategy, in which FLP recombinase would excise a donor molecule and I-SceI would linearise it to stimulate recombination. The full knockout technology did not work because FLP-mediated excision was not detected in the mosquito germline. However, the component tests revealed something more broadly important: I-SceI-induced breaks were repaired predominantly through homology-based pathways rather than simple end joining. We also observed inheritance distortion consistent with both gene conversion and loss of damaged gametes. These results help explain why homing-based systems can work in mosquitoes, while also highlighting why repair pathway choice and gamete viability need to be measured directly in any new drive configuration.

## Introduction

Malaria remains one of the most devastating infectious diseases globally, with 619,000 deaths reported in 2021 alone despite two decades of intensive control effort (WHO, 2022). The persistence of transmission has been attributed to a combination of insecticide resistance in the principal vector Anopheles gambiae, drug resistance in the parasite, and the limitations of current vector control tools (Rodriguez, 2021). Genetic control strategies aim either to suppress vector populations or to modify them so that they no longer transmit pathogens efficiently. Homing-based systems are particularly attractive because the drive mechanism and the genetic load can be coupled: the nuclease both promotes its own inheritance and disrupts a target sequence. In classical homing endonuclease gene (HEG) systems, the endonuclease recognises and cleaves a sequence on the homologous chromosome that lacks the endonuclease cassette. Repair of this double-strand break (DSB) using the endonuclease-containing chromosome as a template converts the cut chromosome into a copy of the drive-bearing allele, producing super-Mendelian inheritance (Burt, 2003).

The same conceptual logic underlies many CRISPR-based synthetic gene drives. In both cases, drive efficiency depends on the outcome of double-strand break repair in the germline. Repair by homology-directed pathways can produce copying of the drive allele, whereas end joining can generate resistant alleles that are no longer recognised by the nuclease. The relative use of homology-dependent repair, non-homologous end joining, microhomology-mediated end joining and other repair pathways is therefore a central determinant of drive performance.

Before CRISPR-based genome editing became routine, gene targeting in insects was pursued using systems that generated linear donor molecules in vivo. A prominent example was the Rong and Golic (2000) strategy in Drosophila, where FLP recombinase releases a targeting molecule from a transgenic donor locus and I-SceI linearises it to expose recombinogenic ends. We attempted to adapt this approach for An. gambiae. The intended design involved three transgenic components: an enzyme line expressing FLP recombinase and I-SceI endonuclease under germline control of the vasa promoter (Papathanos et al., 2009), which drives expression in both male and female germline cells throughout gametogenesis; a donor locus carrying a targeting construct flanked by FRT sites and I-SceI recognition sites; and reporter configurations allowing repair outcomes to be followed genetically and molecularly. I-SceI was chosen as the linearising nuclease because its 18 bp recognition sequence is absent from the An. gambiae genome, precluding off-target cleavage (Windbichler et al., 2007), and because as a natural homing endonuclease it directly models the mechanism of HEG drive.

The full gene-targeting technology was not completed because FLP-mediated excision was not observed in the An. gambiae germline, despite FLP being functional in An. gambiae cell culture. However, the process of testing the component parts generated a detailed set of observations on the consequences of I-SceI cleavage in mosquito germ cells. These observations are arguably more important than the failed targeting platform. They show that I-SceI can be highly active in the An. gambiae germline under vasa promoter control and that the resulting breaks are frequently resolved through homology-associated repair outcomes — encompassing SSA, MMEJ, synthesis-dependent strand annealing (SDSA), and inter-allelic gene conversion — with comparatively little evidence for canonical NHEJ when any flanking homology is available. The experiments also provide evidence that endonuclease cleavage can distort inheritance not only through homing or gene conversion, but also through differential recovery of viable gametes, a finding subsequently corroborated in CRISPR drive systems (Pescod et al., 2024).

Here, we recast those experiments as a repair-outcome study with implications for homing endonuclease and CRISPR-based gene drive systems. Rather than presenting the work primarily as a failed attempt at pre-CRISPR knockout technology, we focus on what the experiments reveal about the repair environment encountered by endonuclease-based drives in An. gambiae, and on what that environment implies for drive efficiency, resistance allele formation, and the interpretation of transmission bias.

## Materials and Methods

### Mosquito strains and rearing

All mosquito strains were based on the An. gambiae G3 wild-type strain (The Gambia). For site-specific integration, the W62 and X6 strains carrying an attP docking site at position 53,037,041 on chromosome 3R were used (Eric Marois, unpublished). Lines were maintained at 28°C, 80% relative humidity under a 12 h:12 h light:dark cycle. Adults were provided 5% glucose solution ad libitum and were blood-fed on mice for egg production.

### Transgenic line construction

The vasaEnzyme (VFS) line was generated by piggyBac-mediated germline transformation with plasmid pXLtTvasEnz, encoding both I-SceI and FLP recombinase under individual vasa promoter/terminator control, with 3xP3-DsRed as a transformation marker. Three independent insertions (VFS1, VFS5, VFS13) were characterised by inverse PCR; all mapped to chromosome 2L within a 1.2 Mb window.

The T2 Reporter line (Windbichler et al., 2011) carries an I-SceI target embedded within a disrupted out-of-frame paxGFP sequence, flanked by paxCFP (sharing pax promoter homology) and actinRFP. The T3 line carries an in-frame paxGFP with an I-SceI site, but lacks the CFP homology present in T2.

The Donor line was created by φC31-mediated integration of pDonorAG5958paxGFPattB into the X6 attP locus; the NewDonor line was integrated into the X1 locus (derived from VFS1) using the same approach, with the transformation marker moved inside the FRT-flanked cassette. Full methodological details available at https://doi.org/10.25560/9864

### Genetic crosses and phenotypic scoring

For I-SceI repair assays, VFS enzyme lines were crossed to the T2 or T3 Reporter line to generate transheterozygotes (VFS/Reporter). These were individually outcrossed to wild-type mosquitoes. F1 progeny were scored for GFP, RFP (actin), and CFP fluorescence under fluorescence microscopy to assign repair class. F1 ratios were tested against neutral Mendelian expectations by χ2 test.

For gamete inviability versus gene conversion assays, VFS5/NewDonor transheterozygotes were crossed to the X1 homozygous line. Non-fluorescent (-/-) F1 were genotyped by PCR to distinguish wildtype amplicons (indicative of gene conversion) from X1-specific amplicons.

### Molecular analysis

Genomic DNA was extracted using the Wizard Genomic DNA Purification Kit (Promega). PCR products amplified across I-SceI target sites were subjected to in vitro I-SceI digest to identify NHEJ events (resistance to digest indicates target modification). PCR products were sequenced (Sanger; Cogenics) with appropriate primers and aligned to reference using ClustalW2. RT-PCR was performed on gonadal tissue to verify enzyme transcription.

## Results

### A transgenic strategy to generate recombinogenic substrates in the mosquito germline

The original objective was to establish targeted gene modification in *An. gambiae* by generating a linear donor molecule in vivo. The system was adapted from the Rong and Golic approach developed in *Drosophila*. A donor construct was designed such that FLP recombinase would excise a molecule flanked by FRT sites, while I-SceI would cleave within the excised molecule to generate free DNA ends adjacent to regions homologous to the target locus. These ends were expected to stimulate homologous recombination with the endogenous target sequence (Supp Figure 1)

**Figure 1.**
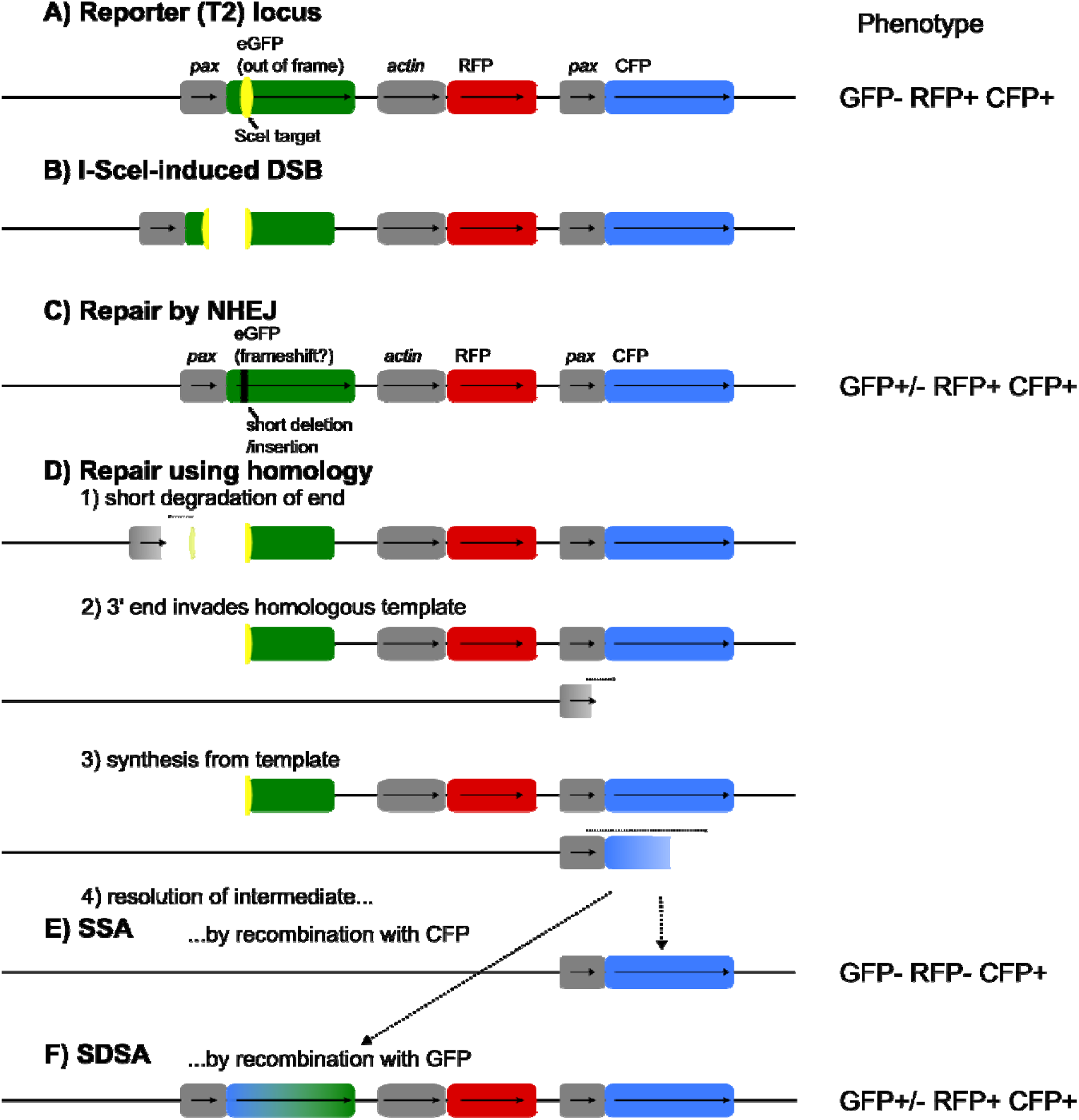
The Reporter line and possible repair outcomes of an I-SceI-induced double strand break (DSB). (A) The Reporter (T2) locus holds the two functional visible markers RFP and CFP. A third marker, GFP, contains an I-SceI target sequence in the coding sequence near the translation start site. This introduces a frameshift into the GFP sequence, preventing translation of functional GFP. (B) I-SceI activity induces a DSB, which may be repaired in different ways. (C) Non-homologous end joining (NHEJ) leads to a loss and/or gain of short sequences, which may result in a framshift that restores the GFP reading frame. (D) Since GFP and CFP have the same 3xP3 (pax) promoter, repair can also take place using this homology. (E) If the resulting repair intermediate is resolved by recombination with CFP, the result (termed Single Strand Annealing, SSA) is the loss of the intervening sequence, including RFP. (F) In Synthesis-Dependent Strand Annealing (SDSA), it can also be resolved by annealing to and recombining with GFP. The phenotype of the original and the repair outcomes are shown on the right.

This design required the independent construction and testing of enzyme, donor and reporter strains. Although the intended donor excision step failed in vivo, the modular design allowed I-SceI activity and downstream repair outcomes to be assayed directly.

### Directing I-SceI activity in the *An. gambiae* germline

A transformation construct was generated in which I-SceI (and FLP) was placed under control of the *vasa* promoter, designed to be active in the mosquito male and female germline (Papathanos et al. 2009) (Supp Fig 2).

**Figure 2.**
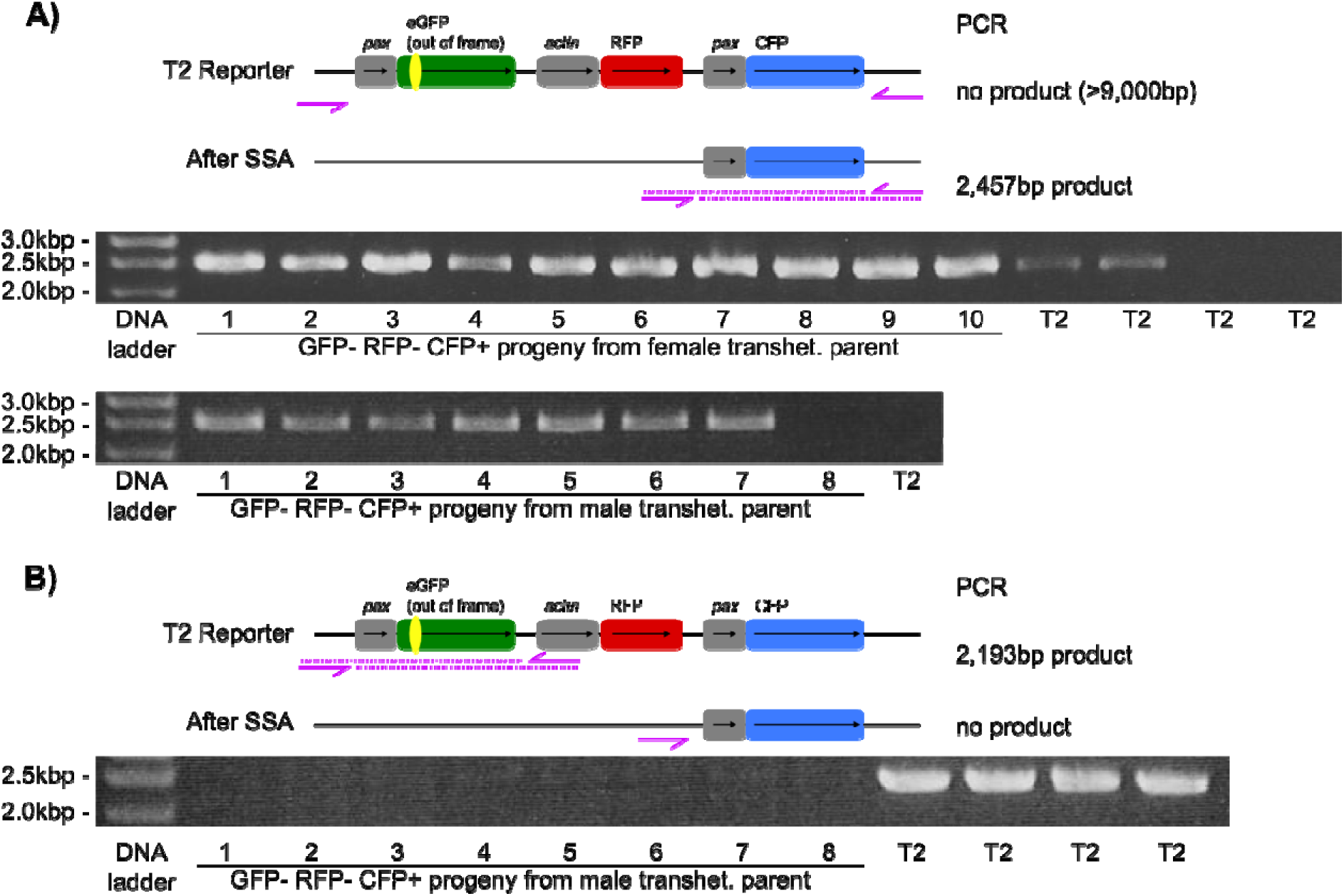
Loss of RFP following Single Strand Annealing. GFP- RFP- CFP+ F1 from male and female enzyme/reporter (*VFS*13/T2) transheterozygote parents were examined for evidence of SSA. Unmodified “T2” Reporter samples were used as controls. (A) The schematic shows the positioning of diagnostic primers (ISceIrepairFor1 and ISceIrepairRev3) on T2 loci before (no product expected) and after SSA (2,457bp amplicon). Amplicons of the size expected for SSA events were obtained. (B) A complementary PCR using primers ISceIrepairFor1 and ISceIrepairRev1 only results in bands in T2 samples, confirming the absence of the full original T2 cassette

Three *VFS* enzyme lines were established by piggyBac-mediated transformation, each containing different single genomic integrations of the same construct, namely *VFS*1, *VFS*5 and *VFS*13.

### I-SceI-induced breaks are preferentially repaired through homology-based pathways

To investigate I-SceI germline activity the VFS enzyme lines were crossed to lines containing a reporter locus (T2) and repair events in transheterozygous parents were recovered in their F1 progeny by fluorescence phenotyping and molecular genotyping. The T2 reporter locus contains a duplicated I-SceI site (one cleavable, one non-cleavable) within an out-of-frame eGFP coding sequence, together with neighbouring fluorescent markers that allowed different classes of repair to be inferred phenotypically (Fig 1A).

Following cleavage by I-SceI the double stranded DNA break (Fig 1B) can be repaired in several different ways. NHEJ or MMEJ resulting in a small insertion or deletion can lead to a frameshift in the non-functional GFP, thus restoring its fluorescence in theoretically 1/3 of such repair events (Fig 1C). The homology between the pax promoters of GFP and CFP can be used for Single Strand Annealing (SSA), leading to a loss of RFP fluorescence (Fig 1D,E). Synthesis-Dependent Strand Annealing (SDSA) is another possible outcome if this homology is used (Fig1F). In SDSA, pax-CFP is used as a template for extension of the broken end, after which the newly-synthesised strand recombines with GFP. As GFP and CFP differ only at a few base pairs, the resulting fluorescent protein may be green or cyan, depending on the crossover point. Hence I-SceI cutting and repair will change the visible markers that are functional. When the enzyme is expressed in the germline, evidence of its activity can be found by examining the fluorescence phenotype of the resulting progeny. Thus offspring with a different phenotype indicate activity of I-SceI in the parental germline. Furthermore, inferences as to the pathway of repair used can be drawn from such phenotypical data and substantiated by genotyping where necessary.

Of 293 F1 carrying the Reporter locus, only 25 (8.5%) retained the parental phenotype, while 243 (82.9%) had lost RFP and 25 (8.5%) had gained GFP (Table 1). This pattern was consistent across all three VFS lines and across both male and female transheterozygous parents, indicating that essentially the entire Reporter locus population had been cleaved and repaired — predominantly by deletion of the intervening sequence (consistent with SSA or MMEJ) or templated synthesis from paxCFP (consistent with SDSA or inter-allelic gene conversion), with no evidence of canonical NHEJ.

**Table 1.** Phenotypic classification of F1 progeny from VFS/T2 Reporter transheterozygote parents.

| I-SceI source | Parent sex | CFP- (WT) | Parental GFP- RFP+ CFP+ | GFP gain GFP+ RFP+ | RFP loss GFP- RFP- CFP+ | Total CFP+ | % Homologous repair |
| --- | --- | --- | --- | --- | --- | --- | --- |
| VFS1 | Male | 57 | 5 | 2 | 38 | 45 | 88.9% |
| VFS1 | Female | 42 | 1 | 3 | 50 | 54 | 98.1% |
| VFS5 | Male | 48 | 7 | 4 | 37 | 48 | 85.4% |
| VFS5 | Female | 52 | 3 | 2 | 39 | 44 | 93.2% |
| VFS13 | Male | 46 | 5 | 8 | 37 | 50 | 90.0% |
| VFS13 | Female | 44 | 4 | 6 | 42 | 52 | 92.3% |
| **Total** | — | **289** | **25** | **25** | **243** | **293** | **91.5%** |

*CFP- F1 lack the Reporter locus. Homologous repair = SSA (RFP loss) + SDSA (GFP gain) as a proportion of all CFP+ F1.*

The overall conclusion is that, at the developmental timing and genomic contexts tested, I-SceI-induced double-strand breaks in the *An. gambiae* germline are frequently channelled into repair pathways that use available homology.

### Molecular characterisation of repair events: SSA or MMEJ, SDSA, and inter-allelic gene conversion dominate; canonical NHEJ absent

To determine the mechanism underlying RFP loss, we performed a diagnostic PCR using primers flanking the entire T2 region. In all RFP- CFP+ samples from VFS13/T2 transheterozygote parents, a band of the predicted deletion product size (2,457 bp) was amplified; no such band was obtained from unmodified T2 controls, confirming that deletion of the intervening sequence between the pax homology regions had occurred.

This is consistent with either SSA using the extended pax promoter homology tracts, or MMEJ exploiting shorter microhomologies within this region; the two mechanisms produce phenotypically indistinguishable outcomes without junction sequencing to identify microhomology signatures at the deletion breakpoints.

GFP gain events are more specifically mechanistically informative: they require templated DNA synthesis rather than simple end deletion, making SDSA or inter-allelic gene conversion the most parsimonious explanations. The observed ratio of parental GFP- to GFP+ was 25:25, significantly deviating from the 2:1 ratio expected if NHEJ frameshift correction were the sole source (χ2, p = 0.012). Sequencing of GFP+ samples confirmed precise deletion of the 47 bp I-SceI-containing insert (Supp Fig 3A), with several samples additionally showing CFP-specific sequence at a dimorphic position 200 bp downstream of the ATG—characteristic of gene conversion tracts extending beyond the DSB, the hallmark of SDSA (Supp Fig 3B).

**Figure 3.**
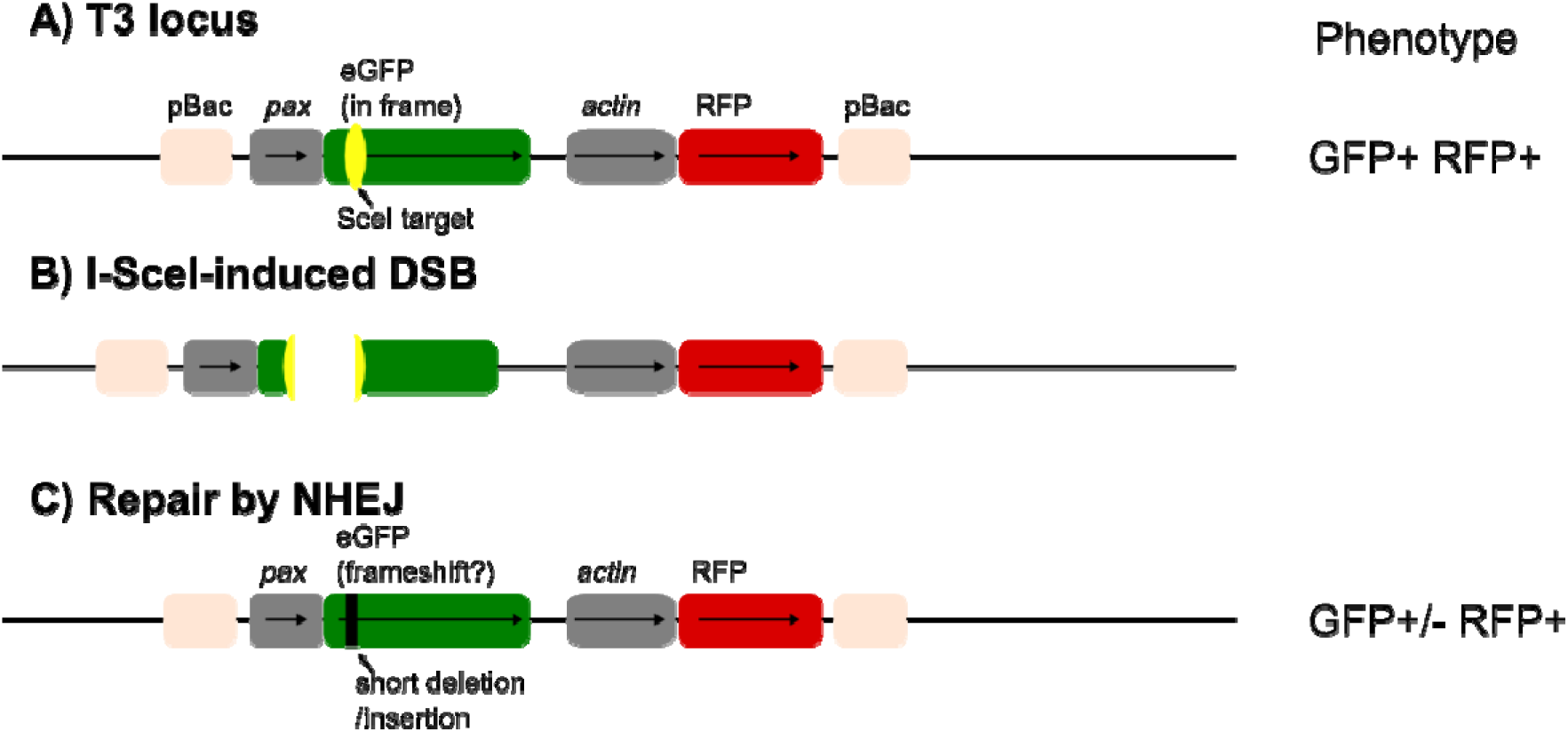
T3 locus, I-SceI cleavage and NHEJ. (A) The T3 locus is similar to the T2 Reporter above, though unlike T2, it lacks CFP and its I-SceI target is in frame, allowing functional eGFP expression. When I-SceI induces a DSB (B), this can be repair by NHEJ (C), which may inactivate the GFP through a frameshift, leading to a GFP- phenotype.

Critically, no canonical NHEJ mutations—random indels at the I-SceI cut site—were recovered in any F1 screened in the presence of flanking homology. This does not exclude perfect end-joining (which would be phenotypically silent) but establishes that the error-prone untemplated joining characteristic of classical NHEJ is effectively suppressed when homology is present. The An. gambiae germline thus appears to resolve DSBs through a hierarchy in which SSA, MMEJ, and SDSA are strongly favoured over canonical NHEJ, consistent with the subsequent finding that even resistance alleles arising in CRISPR drive populations carry MMEJ signatures rather than random indels (Hammond et al., 2017).

### Repair pathway choice is context-sensitive and homology proximity-dependent

To examine repair in a context lacking intrachromosomal homology, VFS1 was crossed to an alternative Reporter line (T3), which contains an in-frame I-SceI site in paxGFP but no paxCFP homology. Instead of the expected ∼50% RFP+ (T3-carrying) F1, only 1 out of 192 scored larvae carried the parental T3 locus (0.5%). No GFP- RFP+ individuals were observed, indicating that NHEJ-induced frameshifting—the expected outcome—was absent. The near-complete disappearance of T3-linked markers implies either (i) gamete inviability from unrepaired DSBs, or (ii) gene conversion from the wildtype allele at that locus, whose native flanking DNA lies only ∼900 bp from the I-SceI site.

#### Evidence for inter-allelic gene conversion and meiotic drive

In the T2 Reporter assay, the ratio of Reporter-carrying (CFP+) to wild-type (CFP-) F1 was 293:289, showing no allelic bias (χ2, p > 0.5). The presence of intrachromosomal homology (paxCFP) allowed efficient repair without allele loss. In contrast, in transheterozygotes where the I-SceI target-bearing locus was one of the donor lines designed for the Rong&Golic gene targeting approach(“ NewDonor”) and lacked internal homology (Supp Fig 4), F1 ratios were consistently skewed: approximately 40% of offspring carried the target allele (versus 50% expected), a significant departure from Mendelian inheritance (χ2, p < 0.0001 in combined analyses; Table 2).

**Figure 4.**
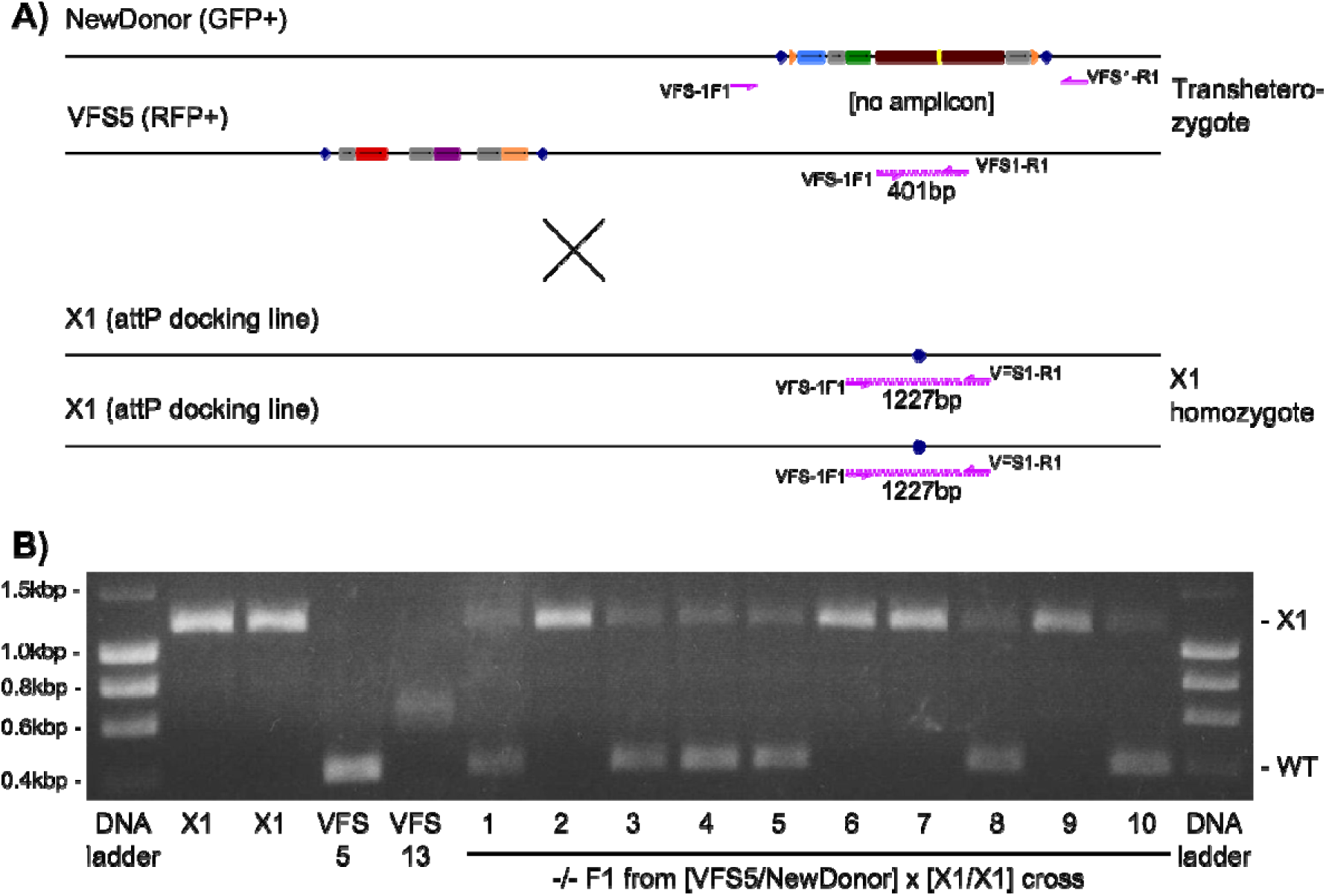
Molecular test of gene conversion in -/- F1. (A) The three different genotypes produce different (or no) amplicons. (B) Outcome of the molecular analysis. Controls X1 and VFS5 show expected amplicons. (Different size of VFS13 amplicon may result from a polymorphism at the locus, but is of no further relevance here.) Ten non-fluorescent (-/-) F1 from a cross of VFS5/NewDonor THs with X1 show presence of wildtype (WT) amplicon in 6 out of 10 samples. (1-3 are F1 from male TH parents, 4-10 originate from female TH parents.) All ten samples show expected band for X1 allele.

**Table 2.**
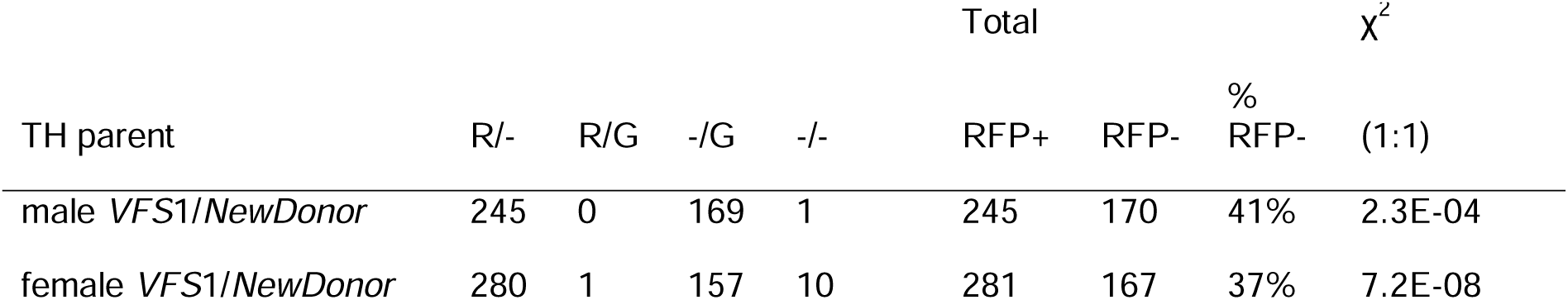
Progeny from VFS1/NewDonor transheterozygous (TH) parents by phenotype. R = RFP+; G = GFP+. %RFP- is the percentage of RFP- of all larvae screened.

To discriminate between gamete inviability (loss of unrepaired chromosomes) and gene conversion (replacement of target allele by the homologous allele), we established transheterozygotes carrying the NewDonor and VFS5 constructs at tightly linked but non-identical loci (0.142 Mb apart; expected recombination rate 0.2%) (Supp Figure 5).

When crossed to the X1 line (Figure 4A), wildtype PCR amplicons—diagnostic of gene conversion—were identified in 6 of 10 non-fluorescent F1 (Figure 4B; Table 2). The remaining 4 could not be explained by gene conversion and likely reflect gamete inviability from unrepaired DSBs. The ratio of VFS5:NewDonor-derived F1 was significantly skewed only in female transheterozygote parents (165:123, p = 0.013), suggesting sex-specific differences in repair outcome consistent with the known biology of male and female An. gambiae germlines (Table 3).

**Table 3.** Test for gene conversion and gamete inviability. Phenotypes of progeny from reciprocal crosses of VFS5/NewDonor transheterozygotes were scored. The p values for the deviation of the RFP+(VFS):RFP-(NewDonor-derived) ratios from 1:1 inheritance are calculated with a χ2 test.

| male parent | female parent | RFP+<br>GFP- | RFP-<br>GFP+ | RFP-<br>GFP- | RFP+:RFP- | $\chi^2$ |
| --- | --- | --- | --- | --- | --- | --- |
| *VFS5/NewDonor*<br>transhet. | X1/X1 | 85 | 72 | 3 | 85:75 | p=0.43 |
| X1/X1 | *VFS5/NewDonor*<br>transhet. | 165 | 79 | 44 | 165:123 | p=0.013 |

### Proximity of homology after chromosome cleavage may alter repair outcomes

The repair outcomes at the NewDonor locus, where the I-SceI site lies ∼5–7 kb from native genomic sequence, contrast with those at the T3 locus, where the equivalent region is shorter at 900bp. At the NewDonor locus, the majority of I-SceI sites remained intact after passage through VFS germlines— likely not because cleavage was absent, but because the lack of nearby homology apparently forced faithful end-rejoining (perfect NHEJ), as predicted by Rong and Golic (2003). Taken together, these data demonstrate that homology proximity to the DSB is a strong determinant of repair pathway in the germline.

### FLP-mediated excision was not detected in the mosquito germline

The gene-targeting strategy depended on FLP-mediated excision of the donor molecule from its chromosomal locus. FLP activity could be demonstrated in *An. gambiae* cell culture using reporter assays, indicating that the enzyme was not intrinsically inactive in anopheline cells. However, crosses designed to detect FRT recombination in the mosquito germline failed to provide molecular evidence for excision at two independent donor loci.

We propose three non-exclusive hypotheses: (i) FLP is transcribed and translated correctly but is insufficiently active at vasa-controlled germline stages due to unfavourable chromatin state or timing; (ii) the proximity of I-SceI and FLP targets (∼4.5–6 kb) leads to antagonistic interference, potentially mediated by γH2AX histone modification spreading ∼3–5 kb from the DSB (Shroff et al., 2004) and overlapping with the FRT sites; (iii) FLP reaction kinetics are fundamentally limiting—as a homotetrameric enzyme requiring four monomers to catalyse recombination, FLP activity scales with the fourth power of concentration, meaning that even modest reductions in expression yield near-zero recombination while I-SceI (monomeric) retains activity.

For the purposes of the present paper, the important point is that the failure of FLP prevented recovery of the intended targeted knockout, but did not prevent assessment of I-SceI cleavage and repair.

## Discussion

The primary finding of this study is that I-SceI-induced double-strand breaks in the *An. gambiae* germline are frequently repaired through homology-dependent pathways. Although the original aim was to establish a pre-CRISPR gene-targeting system based on in vivo donor excision and linearisation, the more durable result is a set of observations on how the mosquito germline responds to endonuclease cleavage.

This has direct relevance to gene drive. Homing-based drives require cleavage of the wild-type allele followed by repair using the drive-bearing homologous chromosome. If the break is repaired by homology-directed copying, drive occurs. If it is repaired by end joining, resistant alleles can be generated. If cleavage causes unrepaired damage or extensive resection, gametes carrying the damaged allele may be lost, producing inheritance distortion that superficially resembles drive but has different population-genetic consequences. Importantly, not all end-joining is equivalent: canonical NHEJ produces unpredictable random indels, while MMEJ uses flanking microhomologies to produce more constrained deletions. When the resistance alleles that arose and were positively selected over 25 generations of CRISPR drive in An. gambiae were sequenced, the dominant class were short in-frame deletions explicable by MMEJ at 3 bp microhomology repeats flanking the cut site — not the random indel spectrum of canonical NHEJ (Hammond et al., 2017). This indicates that even when homing fails and end joining is the only available repair route, the An. gambiae germline preferentially uses homology-sensing mechanisms. Crucially, the same study estimated the meiotic end-joining rate in the germline at only ∼1%, with the majority of resistance alleles arising instead from embryonic end joining caused by maternally deposited Cas9 — a source of resistance that can be substantially reduced by tighter germline-restricted promoters (Hammond et al., 2021).

The data presented here support the view that the An. gambiae germline can be permissive for homology-dependent repair after nuclease cleavage. This is encouraging for homing endonuclease and CRISPR-based drive systems. However, the findings also caution against treating super-Mendelian inheritance as synonymous with accurate homing. Inheritance bias may reflect a composite of true gene conversion, local repair events, marker loss, and selective loss of gametes carrying damaged chromosomes. Distinguishing these outcomes requires molecular assays, not simply phenotypic scoring of progeny. A further practical implication concerns target site selection: because MMEJ deletion endpoints are largely determined by the identity and position of flanking microhomologies, the most probable end-joining repair outcomes at any candidate gRNA or HEG target site can be predicted in silico from the local sequence context. Candidate target sites can therefore be evaluated before drive construction for their likelihood of generating R1 alleles — short in-frame deletions that restore gene function and are strongly selected — versus R2 alleles — frameshift deletions that disrupt function and are not. Prioritising sites where the most probable MMEJ products are frameshifts is a tractable addition to the target site selection workflow, sitting alongside conservation-based criteria and guide RNA efficiency scoring.

Our experiments with the T2 Reporter and donor loci also illuminate the dependency of repair pathway on the proximity of homology to the DSB—a finding with direct implications for real-world drive deployment into wild An. gambiae populations, which harbour extensive natural variation. Two apparently contradictory recent findings frame this discussion. Pescod et al. (2023) found no significant reduction in homing across three geographically diverse An. gambiae strains with 5.3–6.6% target locus heterology (TLH), with homing rates of 81–100% maintained irrespective of strain-of-origin. By contrast, Harvey-Samuel et al. (2026) showed in Culex quinquefasciatus that as little as 6–9% flanking sequence heterology reduced homing efficiency by up to 54%, not by reducing Cas9 cutting efficiency, but specifically by impairing HDR at the repair stage.

Our data offer a mechanistic perspective on why An. gambiae may be more tolerant of flanking sequence heterology than other species. We show that the predominance of HDR is context-dependent: when homology is immediately proximate (T2 Reporter, ∼0 bp from the break to paxCFP homology), essentially all repair is homologous. When homology is more distant (T3 line, ∼900 bp; donor lines, 5–7 kb), outcomes shift towards either gene conversion from the wildtype flanking allele (T3) or near-perfect end-joining of the DSB (donor lines), consistent with data from Rong and Golic (2003) in Drosophila. This proximity dependence suggests that An. gambiae HDR can operate effectively with very short homology tracts—consistent with Pescod et al. (2024) finding that >80% of conversion tracts resolve within 50 bp. If the minimum functional homology for HDR initiation is short in An. gambiae, then moderate levels of sequence divergence in flanking regions may have proportionally less impact on homing efficiency than in species with longer homology requirements.

Beyond homing, we observe evidence for a second mechanism of inheritance bias: gamete-level transmission distortion in the absence of nearby homology (Section 3.4). Where I-SceI cuts a target allele and repair fails or produces gene conversion, the target allele is underrepresented among viable offspring (approximately 40% instead of 50%). Using the VFS5/NewDonor cross design, we demonstrate that this bias arises from two non-exclusive mechanisms: gene conversion of the target allele to the homologous VFS allele, and gamete inviability from unrepaired DSBs. This is mechanistically analogous to meiotic drive, and if the total viable gamete pool is not proportionally reduced—that is, gametes carrying unrepaired chromosomes are simply lost while those from the other allele are not—then the drive-bearing chromosome achieves super-Mendelian transmission independently of homing.

A related strategy was explored by Windbichler et al. (2008), where HEG-mediated cleavage of X-linked ribosomal DNA during spermatogenesis was intended to distort the sex ratio towards males, thereby reducing population reproductive output. That system was limited by over-stability of the nuclease and inadvertent zygotic X chromosome damage, but the underlying logic — that DSB-induced gamete inviability at a target locus can drive inheritance bias — is supported by our observations. More recently, Pescod et al. (2024) demonstrated that both homing and meiotic drive contribute to the super-Mendelian inheritance of CRISPR gene drives in An. gambiae, with >80% of homing conversion tracts resolving within 50 bp of the break. Verkuijl et al. (2022) similarly showed in Aedes aegypti that inheritance bias in a CRISPR drive system could arise through mechanisms other than true HDR homing, reinforcing the point that super-Mendelian inheritance should not be assumed to reflect a single repair mechanism without molecular confirmation. The observations reported here anticipate both findings and provide an early empirical basis for understanding why drive inheritance bias in An. gambiae is likely to be a composite of gene conversion, gamete loss, and repair pathway choice rather than a single clean homing event.

The finding that FLP-mediated excision in the mosquito germline compromised the intended Rong&Golic gene targeting strategy remains technically informative but is not the central biological message. FLP activity in cell culture suggests that the enzyme can function in anopheline cells, but the lack of detectable germline excision at two donor loci indicates that the Rong and Golic strategy cannot simply be transferred from *Drosophila* to *Anopheles* without further optimisation. Possible explanations include chromatin context, timing of enzyme expression, competition between FLP and I-SceI, or kinetic dominance of I-SceI cleavage over FLP recombination. With the advent of programmable nucleases, specifically CRISPR, these limitations are less important as a route to gene targeting, but they remain useful as lessons in the portability of genetic technologies across insects.

These experiments were performed before CRISPR-based gene editing transformed the field of nuclease-based gene drive research. In retrospect, their value lies in the fact that they used I-SceI, a natural homing endonuclease, to interrogate precisely the type of repair decision that determines whether homing-based systems work. The predominance of homology-associated repair outcomes is consistent with the later success of nuclease-based drive approaches in mosquitoes. At the same time, the observation of non-conversion repair and gamete loss anticipates issues that remain central to modern drive design: resistance formation, incomplete homing, sex-specific repair differences and fitness consequences of cleavage.

Overall, the work shows that the *An. gambiae* germline is capable of repairing endonuclease-induced breaks using homologous sequence at high frequency, but that cleavage can also produce complex outcomes beyond simple copying. Future drive designs should therefore measure repair products directly, separate gene conversion from gamete loss, and consider male and female germlines independently.

## Supplementary Figures

**Supp Figure 1.**
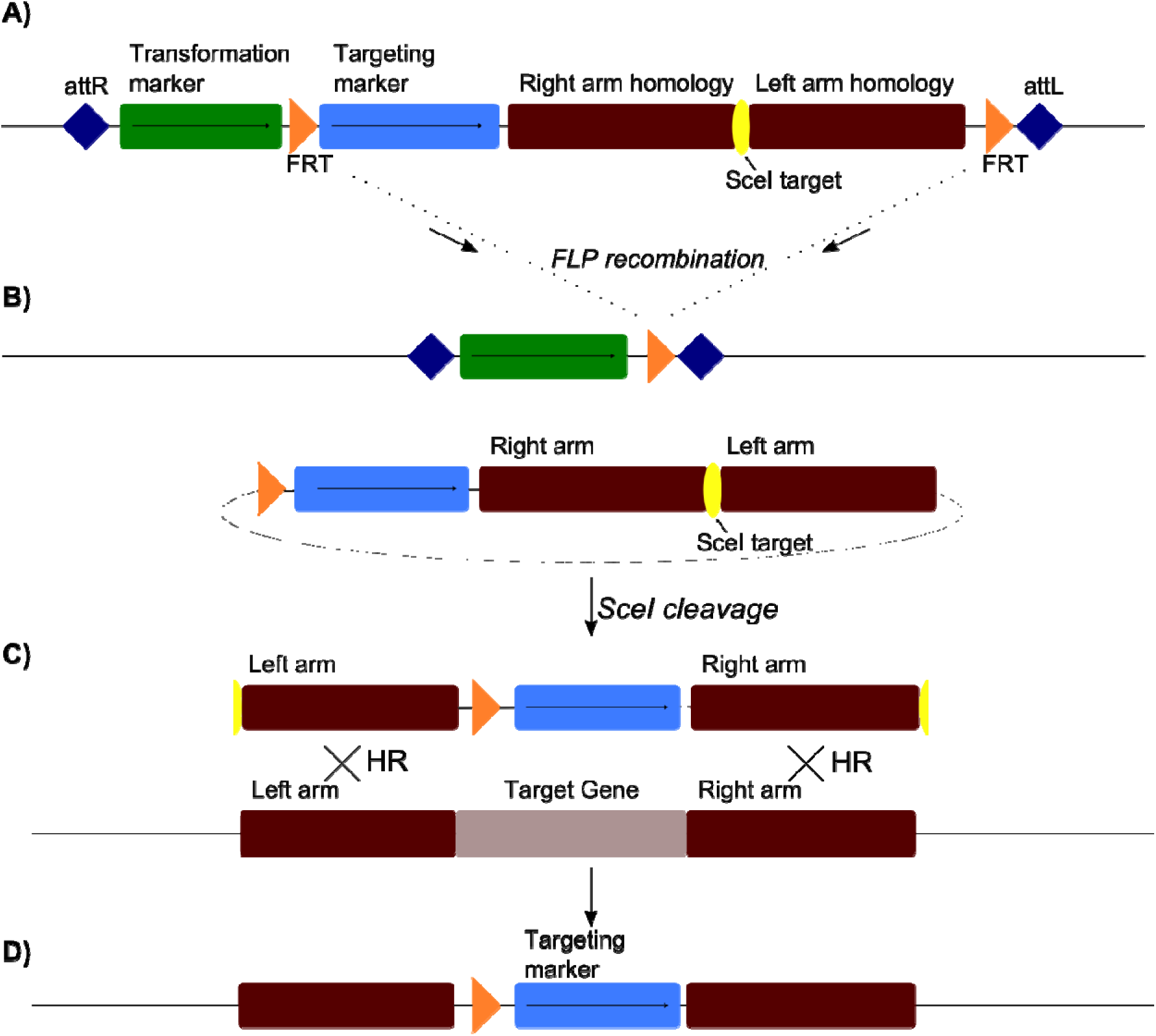
Schematic of gene targeting using Rong & Golic approach. (A) A donor locus containing a marker for transformation and the targeting precursor flanked by two FLP recombination targets, FRT. The targeting precursor a targeting marker and two targeting arms homologous to the targe locus. (B) FLP-mediated recombination excises the circular targeting molecule from the donor locus. (C) I-SceI endonucleases linearises the targeting molecule, which aligns with its target locus. (D) Homologous recombination leads to the integration of the targeting molecule. The targeting marker can be used to recover targeted events.

**Supp Fig 2.**
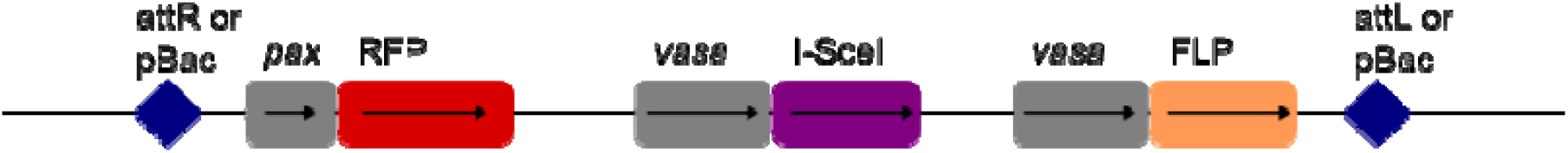
The *VSF* enzyme line. Both I-SceI and FLP enzymes are driven by the *vasa* promoter. *pax*RFP acts as a transformation marker. att sites or piggyBac arms are both shown schematically.

**Supp Figure 3.**
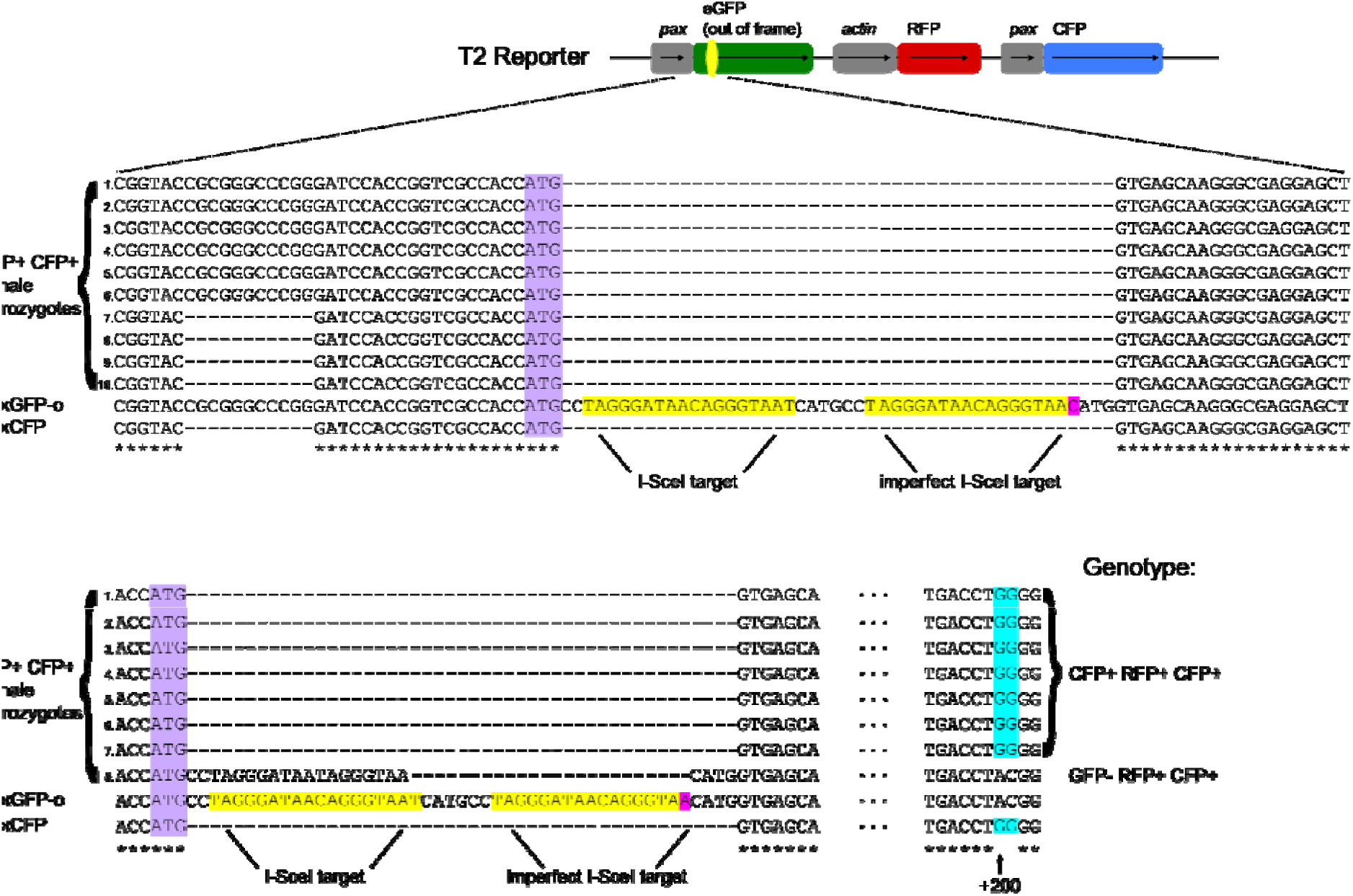
Genotyping of GFP+ and GFP- progeny. GFP+ RFP+ CFP+ and GFP- RFP+ CFP+. F1 samples from *VFS*13/T2 transheterozygote parents were analysed for GFP genotype. An amplicon obtained using ISceIrepairFor1 and ISceIrepairRev1 was sequenced with primer egfp5seq. The results were aligned with both the out-of-frame paxGFP (paxGFP-o) and the functional paxCFP, with the start codon marked in purple. (A) The GFP+ samples have all completely lost the sequence inserted into GFP. paxGFP-o also contains an extra CGCGGGCCCGG in the pax promoter, relative to paxCFP, which was lost in samples 7-10. (B) GFP- samples 1-7 also have the native GFP/CFP sequence directly after the start codon. Further downstream at position +200, the CFP-specific bases GG are found instead AC, as in GFP. Sample 8 has lost the bases between the partially duplicated I-SceI targets.

**Supp Fig 4.**
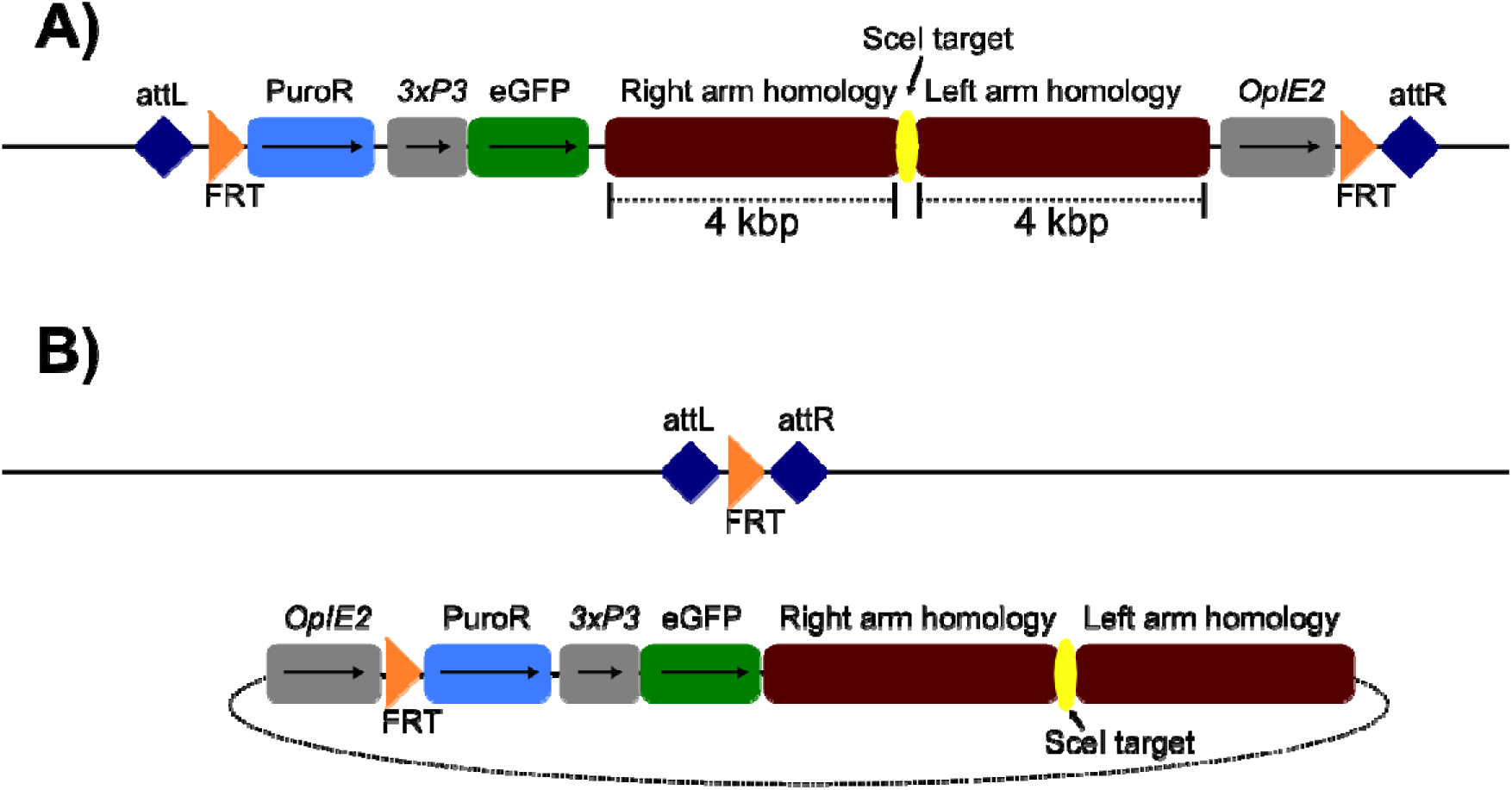
Design of NewDonor cassette. (A) The NewDonor integration will have its transformation marker paxGFP flanked by FRT sites. (B) Should FRT recombination take place, the locus is left markerless, allowing visual identification of FLP activity.

**Supp Figure 5.**
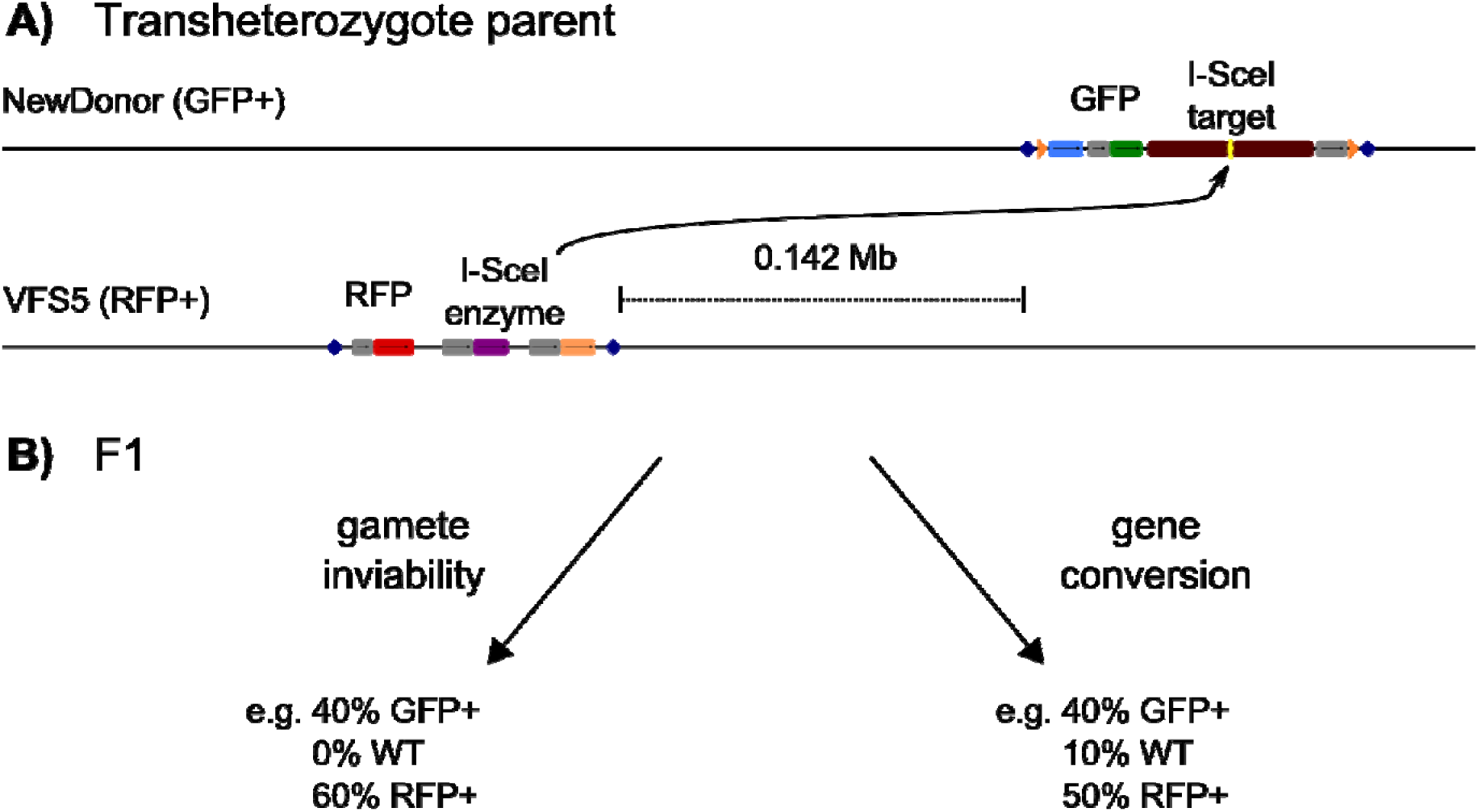
Gamete inviability vs gene conversion crossing scheme. (A) The transheterozygote has the I-SceI target (NewDonor) at a different locus from the I-SceI source (VFS5). Nevertheless, the loci are closely linked (0.142Mb), making recombination unlikely. (B) The two competing hypotheses generate different progeny ratios. Crucially, gene conversion would result in wildtype progeny, while gamete inviability would not. The percentages are merely an example, and the actual percentages may vary, depending on the enzyme source.

